# Fiat lux: A hotspot of *Luciola* fireflies on Western Mediterranean islands

**DOI:** 10.64898/2026.05.27.728198

**Authors:** Andrea Chiocchio, Elisa Serafini, Leonardo Forbicioni, Emiliano Mori, Alessandro Lagrotteria, Leonardo Ancillotto, Andrea Viviano, Roberta Bisconti, Daniele Canestrelli

## Abstract

1. Islands are priority areas for the study and conservation of biodiversity, as they frequently harbour distinct lineages. Yet they are particularly threatened by the global change. In the frame of an accelerated biodiversity crisis, many insular species risk disappearing before they are even discovered.
2. Fireflies, despite their ecological and cultural significance, remain poorly investigated, especially in temperate regions, where they have shown marked decline over the last decades.
3. We investigated firefly diversity across a Mediterranean biodiversity hotspot by genotyping 114 fireflies of the genus Luciola from 29 locations across the Tuscan Archipelago and the adjacent Italian peninsula and applied phylogenetic and species delimitation methods to characterise genetic differentiation and its geographic structure.
4. We found an unexpectedly high level of genetic differentiation in this area. Phylogenetic inference recovered six deeply divergent and geographically structured mitochondrial lineages: three restricted to Elba Island and three to the Italian Peninsula. Genetic divergence among these lineages ranged from 1.97% to 4.58%, values comparable to or exceeding those typically observed among distinct species. Accordingly, species delimitation methods consistently supported their status as distinct species.
5. The coexistence of three divergent lineages on Elba Island suggests a biogeographic scenario characterised by ancient island colonisations and possible in situ diversification. These findings reveal a previously unrecognised depth of evolutionary diversity in Italian fireflies and identify the Tuscan Archipelago as a priority area for future research on firefly evolution and conservation, emphasizing that fireflies are a major gap in our knowledge of insect biodiversity in Europe.

## Introduction

In the context of an accelerating biodiversity crisis, scientists are increasingly concerned that many species may be lost before they are even discovered (Costello et al., 2013; Pereira et al., 2010; Stork, 2010). This concern stems from the recognition that the most speciose taxa remain severely understudied and that the most species-rich regions of the planet remain underexplored, despite being profoundly affected by global change. Furthermore, recent integration of field-based research and museum inventories with genetic investigations have uncovered substantial cryptic biodiversity in virtually all taxa, even in well-studied geographic areas. This evidence prompted conservation initiatives aimed at halting biodiversity decline, while simultaneously intensifying efforts to document biodiversity before it is too late.

Islands often host high levels of biodiversity and constitute ecologically unique environments yet contain some of the most vulnerable ecosystems (Whittaker & Fernández-Palacios, 2007; Losos & Ricklefs, 2009; Whittaker et al., 2017). Because of the peculiar environmental conditions and geographic isolation, insular populations often show phenotypic and/or genetic differentiation from mainland populations. For this reason, island biodiversity is typically represented by endemic units, and their extinction would result in the loss of irreplaceable biological diversity (Kier et al., 2009; Losos & Ricklefs, 2009). Furthermore, the limited distributions and small population sizes make island species extremely vulnerable to the major drivers of biodiversity decline, such as the introduction of alien species, land-use change, and climate change (Wood et al., 2017). Nevertheless, despite the strong efforts devoted to the study and conservation of island biodiversity, unexpected new species continue to be discovered, even among ecologically significant and charismatic taxa (Xin et al., 2025)

Fireflies (Coleoptera: Lampyridae) are fascinating worldwide-distributed insects, that use light spots as mating signals to attract partners at night (Lewis et al., 2020). Historically valued as natural light sources and deeply embedded in cultural traditions, fireflies attract today considerable attention as bioindicators of land use changes and light pollution (Napompeth, 2009; Lewis, 2016; Ghosh et al., 2024). A growing number of observations report that firefly populations have been declining in recent decades across many regions, primarily due to light pollution, which interferes with their bioluminescent mating communication, but also due to habitat loss, population fragmentation and climate change (Fallon et al., 2021; McNeil et al., 2024; Picchi et al., 2013; Chatragadda, 2020; Mbugua et al., 2020). In spite of the documented decline, fireflies remain underrepresented in global conservation priorities, largely due to major knowledge gaps in their taxonomy, distribution, and ecology (Lewis et al., 2020). Indeed, recent taxonomic and systematic investigations are revealing a large number of unrecognised cryptic lineages, especially in tropical and subtropical regions (Jusoh et al., 2020; Jusoh et al., 2021). At the same time, integrative taxonomic studies combining morphological and molecular phylogenetic approaches have revealed marked genetic differentiation and several cryptic species also within European fireflies (Mori et al., 2025; Nunes et al., 2026). This evidence suggest that species diversity in temperate fireflies may be higher than currently recognised, highlighting the need for further investigation (Lagrotteria et al., 2026; Lewis et al., 2024; Nunes et al., 2026).

In this study, we investigated firefly species diversity in one of the biodiversity hotspots of the Mediterranean region, the Tuscan Archipelago (Mittermeier et al., 2005; Ruzzier et al., 2021). Located in the northern Tyrrhenian Sea, between the Italian Peninsula and Corsica island, the Tuscan Archipelago acted as an ecological and biogeographic corridor between Corsican and central Italian biotas during the Plio-Pleistocene, when climate-driven sea-level fluctuations created temporary connections among them (Van Andel & Tzedakis 1996; Rohling et al.1998). Consequently, this area harbours a surprisingly rich biota, including numerous endemic and sub-endemic taxa and genetic lineages of both Tyrrhenian island and mainland origin (Dapporto & Cini, 2007; Vignoli et al., 2007; Fattorini, 2010; Fattorini & Dapporto, 2014; Dapporto et al., 2017; Ancillotto et al., 2026; Lagrotteria et al., 2026). Notably, despite the large number of studies on the flora and fauna of this region, firefly diversity remains poorly characterised. A recent survey reported three fireflies species living within the Tuscan archipelago, i.e. *Lampyris plurihomonyma, Lampyris fuscata* and *Luciola pedemontana* (Lagrotteria et al., 2026). Amongst those, *Luciola pedemontana* resulted the most widespread, being recorded in at least three islands (Elba, Giglio, and Pianosa) with high population densities. In this work, we conducted extensive sampling of *L. pedemontana* populations across the Tuscan Archipelago and used molecular markers to assess their genetic identity and genetic structure in comparison with populations from the nearby Italian Peninsula.

## MATERIALS AND METHODS

Between 2021 and 2024, we collected 114 firefly individuals from 29 sites distributed across the Tuscan Archipelago and the Italian peninsula (Figure 1; Table 1). All individuals were preliminarily identified and assigned to *Luciola pedemontana* ([Curtis] 1843) based on morphological features, that is a red pronotum with no black spots (Fanti, 2022). Specimens were collected manually or using a hand net, with a maximum of 1 to 5 individuals per site to minimise potential impacts on local populations, and stored singularly in 96% ethanol until DNA extraction. Insect collection procedures followed the relevant guidelines and have been approved by the Tuscan Archipelago National Park (permit number 9631/2017).

**Table 1.** Geographical location of the 29 collection sites of *Luciola* specimens analysed in this study, with sample size (n) and specimen code.

| Species | Site Code | Locality | Latitude (N) | Longitude (E) | n | Specimen ID |
| --- | --- | --- | --- | --- | --- | --- |
| *Luciola pedemontana* | 1 | Casentino Forests-Camaldoli | 43.784 | 11.881 | 6 | LUCAS02, LUCAS03_S, LUCAS05_S, LUCAS04_S, LUCAS06, LUCAS07 |
|  | 2 | Tarquini-Borgo Saline | 42.202 | 11.722 | 13 | LUCTAR1, LUCTAR2, LUCTAR3_S, LUCTAR4, LUCTAR05_S, LUCTAR6, LUCTAR07_S, LUCTAR8, LUCTAR09_S, LUCTAR10, LUCTAR11, LUCTAR12, LUCTAR13 |
|  | 3 | PNALM- Val Fondillo | 41.779 | 13.857 | 3 | LUCABR01_S, LUCABR02_S, LUCABR03_S |
|  | 4 | Catanzaro-Nocella Serrastretta | 39.013 | 16.417 | 1 | LUCAL3_S |
|  | 5 | Ponza Island-Punta Incenso | 40.931 | 12.983 | 8 | LUCPON7_S, LUCPON6, LUCPON9_S, LUCPON4, LUCPON8, LUCPON11_S, LUCPON10_S, LUCPON21 |
|  | 6 | Ponza Island-Port | 40.893 | 12.962 | 13 | LUCPON14, LUCPON1, LUCPON17, LUCPON13, LUCPON5, LUCPON3_S, LUCPON19, LUCPON22, LUCPON16, LUCPON20, LUCPON15, LUCPON02_S, LUCPON18 |
|  | 7 | Elba island-Pomonte | 42.749 | 10.125 | 7 | LUCELB21.1, LUCELB21.2, LUCELB28.2, LUCELB28.3, LUCELB21.4, LUCELB21.3, LUCELB28.1 |
|  | 8 | Elba island-Le secche | 42.822 | 10.371 | 3 | LUCELB18.1, LUCELB18.2, LUCELB18.3 |
|  | 9 | Elba island-Nisportino | 42.823 | 10.383 | 6 | LUCELB17.1, LUCELB17.2, LUCELB14.1_S, LUCELB14.2_S, LUCELB14.3_S, LUCELB14.4 |
|  | 10 | Elba island-Mount Serra | 42.831 | 10.409 | 3 | LUCELB19.1_S, LUCELB19.2_S, LUCELB19.3 |
|  | 11 | Elba island-Mount Capanne | 42.786 | 10.167 | 1 | LUCELB20.4_S |
|  | 12 | Elba island-Cavo | 42.856 | 10.426 | 4 | LUCELB23.1, LUCELB23.2_S, LUCELB23.3, LUCELB23.4 |
|  | 13 | Elba island-Marciana | 42.789 | 10.172 | 2 | LUCELB25.2, LUCELB25.3 |
|  | 14 | Elba island-Campo al Pero | 42.797 | 10.403 | 1 | LUCELB26.1_S |
|  | 15 | Elba island-Rio Marina | 42.813 | 10.426 | 3 | LUCELB27.1_S, LUCELB27.2, LUCELB27.3_S |
|  | 16 | Elba island-Norsi | 42.770 | 10.343 | 2 | LUCELB3.1_S, LUCELB3.2_S |
|  | 17 | Elba island-Botro | 42.776 | 10.385 | 3 | LUCELB9.1_S, LUCELB9.2_S, LUCELB9.3_S |
|  | 18 | Elba island-Mola | 42.759 | 10.385 | 3 | LUCELB1.1_S, LUCELB1.2_S, LUCELB1.3 |
|  | 19 | Elba island-Mount Orello | 42.784 | 10.327 | 2 | LUCELB11.02, LUCELB11.3_S |
|  | 20 | Elba island-Acquaviva | 42.819 | 10.287 | 3 | LUCELB6.1, LUCELB6.2_S, LUCELB6.3 |
|  | 21 | Elba island-Nisporto | 42.824 | 10.381 | 3 | LUCELB13.1_S, LUCELB13.2_S, LUCELB13.3_S |
|  | 22 | Elba island-San Martino | 42.785 | 10.290 | 3 | LUCELB15.1, LUCELB15.2, LUCELB15.3_S |
|  | 23 | Elba island-Sant'Andrea | 42.803 | 10.139 | 3 | LUCELB7.1_S, LUCELB7.2, LUCELB7.3_S |
|  | 24 | Elba island-Via del Lentisco | 42.747 | 10.220 | 2 | LUCELB8.1, LUCELB8.3_S |
|  | 25 | Elba island-Colle Reciso | 42.783 | 10.314 | 4 | LUCELB12.02, LUCELB12.03, LUCELB12.04_S |
|  | 26 | Elba island-Gualdo | 42.750 | 10.387 | 4 | LUCELB2.1, LUCELB2.2, LUCELB2.3, LUCELB2.4 |
|  | 27 | Elba island-Acquabona | 42.786 | 10.346 | 3 | LUCELB4.1_S, LUCELB4.2_S, LUCELB4.3 |
|  | 28 | Elba island-San Giovanni | 42.797 | 10.321 | 4 | LUCELB5.1, LUCELB5.2, LUCELB5.3, LUCELB5.4_S |
| *Luciola Italica* | 29 | Giglio Island | 39.940 | 11.646 | 1 | LUCG01_S |

**Figure 1.**
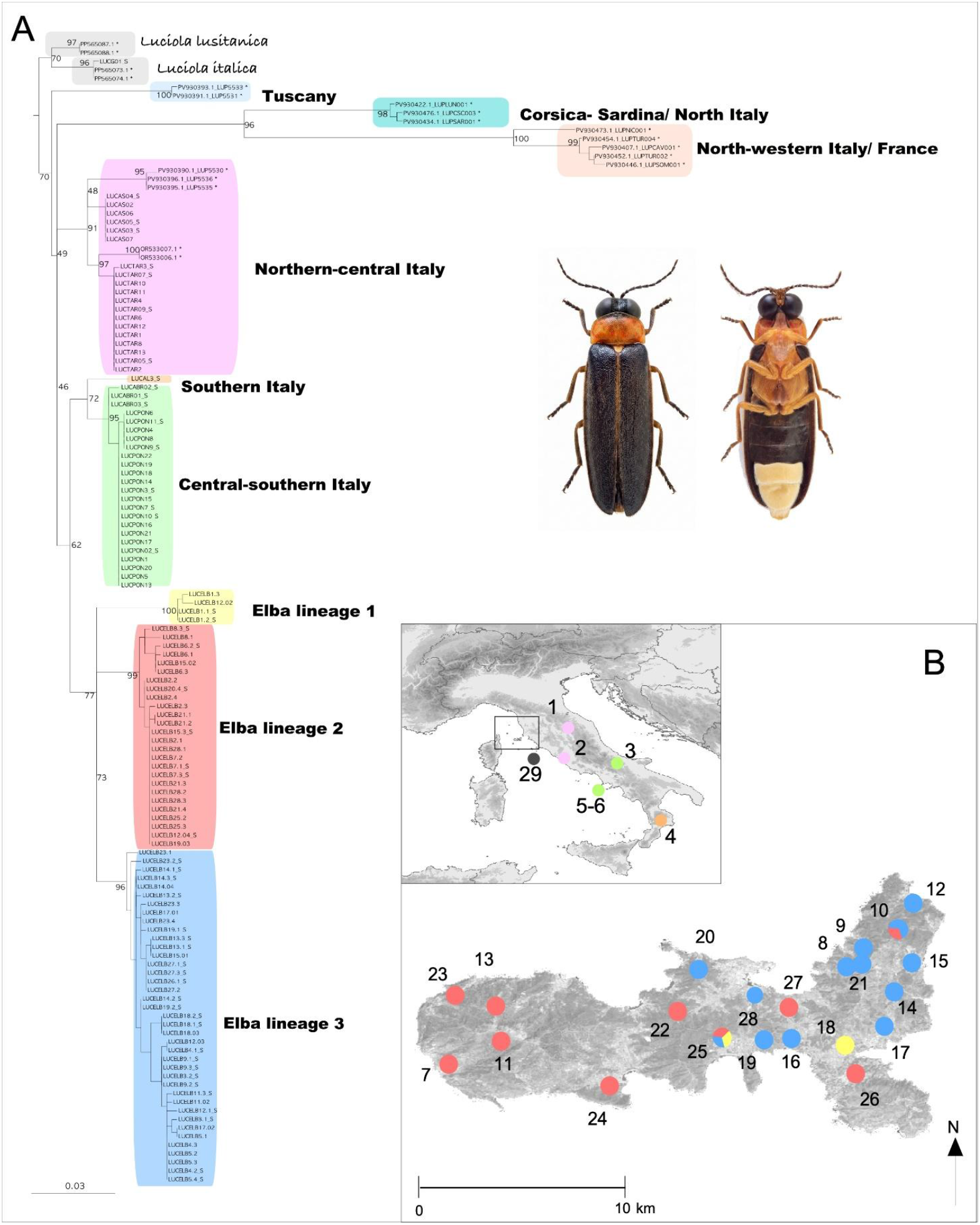
Phylogenetic relationships and geographic distribution of the *Luciola* specimens analysed in this study. (A) Maximum-likelihood consensus tree inferred in IQ-TREE from mitochondrial Cytochrome Oxidase I (COI) gene sequences: branch support values (%) are reported at relevant nodes; colours indicate well-supported distinct lineages; asterisks denote reference sequences obtained from the literature. (B) Geographic distribution of the 29 sampling sites analysed in this study: collection sites are numbered as in Table 1, and colours are associated to the lineages highlighted in the phylogenetic tree.

### DNA extraction and genotyping

Two to four legs were used for DNA extraction. To enhance DNA extraction yield, these tissue portions were previously mechanically ground using a FastPrep® tissue homogenizer (MP Biomedicals) and Lysing Matrix A. DNA was then extracted using Quick-DNA Miniprep Plus Kit (Zymo Research, Irvine, CA), following the manufacturer’s recommendations.

We amplified and sequenced the 658 bp “barcode” fragment of the mitochondrial gene cytochrome oxidase subunit I (COI) using the Oxford Nanopore MinION platform (Oxford Nanopore Technologies, Oxford, UK). Amplifications were carried out by PCR using the universal primers LCO1490 and HCO2198 (Folmer et al., 1994), in a 14 μL reaction mix which also included 20-40 μg of genomic DNA, buffer 5 ×, 3.75 mM MgCl2, 0.3 μM dNTPs, 0.3 μM of each primer, and 0.5 U of Taq polymerase (©Promega Corporation, Madison, Wisconsin, USA). PCR conditions included an initial denaturation at 94°C for 1 min, followed by 35 cycles of 94 °C for 30′′, annealing at 52°C for 1 min, extension at 72 °C for 1 min, and a final extension at 72 °C for 10 min. PCR products were run on 1% agarose gels by electrophoresis to determine whether an amplicon had been obtained for each PCR reaction. Amplicons from each individual were tagged in a further single PCR step using primers with a unique combination of 13-bp sequences to enable high-throughput multiplexing within a single sequencing run (Srivathsan et al., 2019). Afterwards, PCR products were pooled and purified using Sera-Mag beads (GE Healthcare Life Sciences, Marlborough, Massachusetts, USA) according to the manufacturer’s recommendations. Before library preparation, the purified product was quantified using a Qubit™ Flex Fluorometer with a dsDNA assay kit (Thermo Fisher Scientific, Waltham Massachusetts, USA).

MinION sequencing libraries were prepared using the SQK-LSK114 ligation sequencing kit (Oxford Nanopore Technologies) as per the manufacturer’s instructions, starting with 200 ng of pooled PCR product. Except for omitting the FFPE DNA repair mix during the end-repair step, since it is primarily for formalin-fixed, paraffin-embedded samples, the protocol was followed. The final pool was sequenced on a MinION sequencer (Mk1B; Oxford Nanopore Technologies) with an R10.4.1 flow cell. Reaction volumes for libraries were 47 μl DNA, 3.5 μl Ultra II End-prep reaction buffer (New England Biolabs, Ipswich, Massachusetts, USA), 3 μl Ultra II End Prep enzyme mix (New England Biolabs), and 6.5 μl molecular-grade water. The sequencing run was initiated in MinKNOW (v24.06.16; ONT) with starting flow cell pore availability at 1.550 and lasted 20 h.

### MinION data processing

Raw reads were basecalled with Guppy (v6.5.7; ONT; high accuracy base calling mode; parameters: -qscore_filtering --compress_fastq -c dna_r10.4.1_450bps_hac.cfg --min_qscore 7). Demultiplexing, quality filtering, and consensus barcode calling were performed using ONTbarcoder (Srivathsan et al., 2021), a pipeline specifically designed for Nanopore-based DNA barcoding. ONTbarcoder was run in standard (post-run) mode. The software automatically trims primer regions, demultiplexes reads based on specific tags, and generates consensus barcodes using an iterative alignment-based approach. For quality filtering, default ONTbarcoder parameters were used, except for tag-mismatch tolerance and minimum read length, which were set to allow up to 2 bp mismatches from the expected tag sequence and to retain reads ≥450 bp, respectively. Consensus sequences were first generated using ONTbarcoder and subsequently polished with Medaka (v2.0.1; Oxford Nanopore Technologies Ltd., UK), executed with default settings, to further improve base-level accuracy using a neural network-based error-correction model trained on Nanopore data. Afterwards, the polished reads were aligned using MAFFT v7 (Katoh et al., 2013).

As a validation procedure, a subset of specimens was sequenced twice using the Sanger method, using the same PCR products employed for MinION sequencing. PCR product purification and sequencing were carried out by Macrogen Europe (Milan, Italy); the resulting chromatograms were visually inspected using FinchTV v1.4.0. The DNA sequences obtained from Sanger sequencing were then aligned using AliView v1.30 (Larsson, 2014) and compared with those from MinION sequencing to assess consistency.

### Sequence data analysis and species identification

We assessed the genetic identity of the sequenced individuals using a phylogenetic comparative framework. First, we retrieved additional COI sequences from the NCBI database representing other *Luciola* species inhabiting Europe: two sequences of *L. italica* (Linnaeus, 1758) from Sud Tyrol (northern Italy), two sequences of *L. lusitanica* (Charpentier, 1825) from Portugal, and fifteen sequences of *L. pedemontana* representative of the main known lineages (Supplementary Table 1). These sequences were used as references for species identification. Then, we estimated phylogenetic relationships between the reference sequences and our specimen sequences using the maximum likelihood (ML) method implemented in IQTREE v2.4.0 (Minh et al., 2020). The best-fit substitution model was determined using the ModelFinder function implemented in IQ-TREE (Kalyaanamoorthy et al., 2017). Branch support was evaluated using 1000 replicates of the Ultrafast Bootstrap Approximation (UFB; Hoang et al., 2018). Based on the clades identified in the phylogenetic tree, we calculated average genetic distances both within and between clades using the Kimura 2-parameter model implemented in MEGA v11.0.13 (Kimura 1980; Tamura et al. 2021). We also estimated the pairwise genetic distances between individual sequences using the same settings.

Finally, we evaluated the occurrence of multiple putative species within our dataset by means of the most used DNA-based species delimitation tools: the Automatic Barcode Gap Discovery (ABGD; Puillandre et al., 2012), the Bayesian implementation of the Poisson Tree Processes (bPTP; Zhang et al., 2013), and the General Mixed Yule Coalescent model (GMYC; Pons et al., 2006). For this stage, we integrated sequences from all species of the genus *Luciola* available in public databases, after filtering for data quality (Supplementary Table 1). This integration enabled a more robust framework for species delimitation, allowing a broader representation of the genetic differentiation within this species group. Automatic Barcode Gap Discovery (ABGD) was conducted on the “Full_dataset” (see Supplementary material) using the ABGD webserver (https://bioinfo.mnhn.fr/abi/public/abgd/abgdweb.html; accessed on April 5, 2025) with default parameters (*i*.*e*., using Jukes-Cantor model (JC69) distances, a relative gap width of 1 and 50 steps, Pmin = 0.001, Pmax = 0.1, and Nb bins for distance distribution = 20) (Puillandre et al., 2012).

For the GMYC and bPTP analyses, we used the “Subsample_dataset” (see Supplementary material), consisting of a representative subsample that captures the variability within and among the main clades. This choice was motivated by previous evidence that a high number of haplotypes can decrease the efficiency of tree-based species delimitation methods, since oversampling increases the probability of detecting intermediate haplotypes between closely related species (Meyer & Paulay 2005; Puillandre et al. 2012; Phillips et al. 2019; Magoga et al. 2021). The phylogenetic trees used for GMYC and bPTP analyses were generated through the Bayesian approach implemented in BEAST 2.7.6 (Drummond et al. 2012). We employed the Yule pure-birth speciation model as the tree prior, alongside a strict molecular clock model and the most appropriate molecular evolution model determined using Jmodeltest 2.1.10 (Darriba et al. 2012), based on the Bayesian Information Criterion. The Monte Carlo Markov Chain (MCMC) analysis ran for 100 million generations, with trees being sampled every 10000 generations. To assess the independence of the effective sample size (ESS), we ensured that ESS values exceeded 200 for all parameters, using Tracer 1.6, after discarding the first 10% of the samples as burn-in. The consensus tree was then constructed in TreeAnnotator 2.7.6 (part of the BEAST package) by applying the maximum clade credibility criterion, with the first 1000 sampled trees discarded as burn-in. The single- and multiple-threshold GMYC analyses were conducted in R v4.2.0 on a Windows platform using the *splits* package (Ezard et al., 2015). The bPTP analysis was performed on the web-based platform (https://species.h-its.org/ptp/), with an MCMC length of 500000 generations and a 20% burn-in.

## RESULTS

We successfully amplified and sequenced a 648-bp fragment of the mitochondrial COI gene from 114 *Luciola* samples. No insertions, deletions, stop codons, or nonsense mutations were observed. We identified 41 distinct haplotypes, defined by 74 (12.7%) variable positions. The mean haplotype diversity (Hd) and nucleotide diversity (π) values for this dataset were 0.947 (± 0.010 SD) and 0.028 (± 0.001 SD), respectively. The average number of nucleotide differences among sequences was 16.4. All sequences generated in this study, along with their corresponding GenBank accession numbers, are listed in Supplementary Table 2. The ML tree of the mtDNA sequences obtained by IQTREE (log-likelihood score = −2106.398; s.e. = 118.6040) clearly defined one clade for *L. lusitanica*, one clade for *L. italica*, and one clade for *L. pedemontana*, with strong bootstrap support (> 85). No specimens clustered with *L. lusitanica*, and only one specimen from Giglio island clustered with *L. italica* (99,83% identity with reference sequences). All the other specimens clustered with *L. pedemontana*. However, from the inspection of the ML tree, we identified at least 5 distinct and well differentiated genetic lineages among our sequences of *L. pedemontana*: one lineage included samples from northern-central Italy, a second lineage included samples from central and southern Italy, with further substantial differentiation between the samples from central–southern and southern Italy, and three further lineages resulted exclusive of the Elba Island (Fig. 2). Moreover, our phylogenetic reconstruction identified three further lineages among the reference sequences that were markedly differentiated from all the other *L. pedemontana* lineages, in line with what was reported by Nunes et al. (2026): one distributed across Corsica, Sardinia and northern Italy, one restricted to north-western Italy and southern France, and one additional lineage occurring in Tuscany.

The pairwise genetic distances among the lineages identified in the phylogenetic tree are provided in Table 2. The genetic distance between *L. italica* and *L. lusitanica* is 2.15%, while the genetic distance between the identified lineages of *L. pedemontana* and the other two sister species ranged from 3.35% to 4.56% and from 2.33% to 3.42%, respectively. The genetic distance between the lineages identified within *L. pedemontana* ranged from 1.97% to 4.54%, with intra-clade divergence ranging from 0.002 to 0.013. Distances between the mainland and Elba Island clades range from 3.4% (southern Italian lineage vs Elba lineage 1) to 4.58% (central-northern Italian lineage vs Elba lineage 1), and even 10% if consider the northwestern Italy + France clade from the reference sequences. Finally, the genetic distance between the Elba clades and the Sardinia-Corsica clade ranged from 8.1% and 8.9%.

**Table 2.** Reduced matrix of pairwise genetic distance between and within the lineages of *Luciola pedemontana* and the sister species *L. lusitanica* and *L. italica*, as estimated in MEGA (details in the text): values below the diagonal represent inter-clade distances, with standard error among square brackets; bold values on the diagonal indicate mean intra-clade distances, with standard error among square brackets.

|  | Elba<br>lineage_1 | Elba<br>lineage_3 | Elba<br>lineage_2 | southern<br>Italy | central-southern<br>Italy | northern-central Italy | Cosica_Sardi-<br>nia / North<br>Italy | north-western Italy<br>/ France | Tuscany | Luciola<br>italica | Luciola<br>lusitanica |
| --- | --- | --- | --- | --- | --- | --- | --- | --- | --- | --- | --- |
| Elba lineage_1 | **0.004** |  |  |  |  |  |  |  |  |  |  |
| Elba lineage_3 | 0.041<br>[0.008] | **0.006** |  |  |  |  |  |  |  |  |  |
| Elba lineage_2 | 0.040<br>[0.008] | 0.030<br>[0.007] | **0.008** |  |  |  |  |  |  |  |  |
| southern Italy | 0.034<br>[0.008] | 0.041<br>[0.009] | 0.040<br>[0.008] | - |  |  |  |  |  |  |  |
| central-southern Italy | 0.040<br>[0.008] | 0.039<br>[0.008] | 0.037<br>[0.008] | 0.020<br>[0.006] | **0.002** |  |  |  |  |  |  |
| northern-central Italy | 0.046<br>[0.009] | 0.038<br>[0.007] | 0.044<br>[0.008] | 0.035<br>[0.007] | 0.029<br>[0.006] | **0.012** |  |  |  |  |  |
| Cosica_Sardinia_North<br>Italy | 0.090<br>[0.013] | 0.081<br>[0.013] | 0.084<br>[0.013] | 0.079<br>[0.013] | 0.081<br>[0.013] | 0.072<br>[0.011] | **0.004** |  |  |  |  |
| north-western Italy /<br>France | 0.100<br>[0.013] | 0.092<br>[0.013] | 0.096<br>[0.013] | 0.088<br>[0.013] | 0.093<br>[0.013] | 0.087<br>[0.012] | 0.078<br>[0.011] | **0.018** |  |  |  |
| Tuscany | 0.057<br>[0.010] | 0.048<br>[0.009] | 0.046<br>[0.009] | 0.049<br>[0.010] | 0.048<br>[0.009] | 0.040<br>[0.008] | 0.081<br>[0.012] | 0.075<br>[0.011] | **0.002** |  |  |
| Luciola italica | 0.046<br>[0.009] | 0.034<br>[0.007] | 0.042<br>[0.008] | 0.040<br>[0.009] | 0.035<br>[0.008] | 0.034<br>[0.007] | 0.012<br>[0.073] | 0.012<br>[0.084] | 0.009<br>[0.044] | **0.001** |  |
| Luciola lusitanica | 0.032<br>[0.007] | 0.033<br>[0.007] | 0.034<br>[0.008] | 0.032<br>[0.007] | 0.023<br>[0.006] | 0.031<br>[0.007] | 0.012<br>[0.077] | 0.012<br>[0.084] | 0.009<br>[0.042] | 0.021<br>[0.006] | - |

The results from the species delimitation analyses are summarised in Table 3. The ABGD analysis revealed a significant gap in the pairwise distance distribution between 0.06 and 0.07. By applying the prior maximal distance (Pmax) threshold within this range, ABGD generated a primary partition that identified thirty-six groups within the genus *Luciola* (Supplementary Table 3). Focusing on our specimens, *L. pedemontana* was subdivided into six distinct putative species, consistent with the distinct lineages identified in the ML phylogenetic tree: three from the Island of Elba, one from northern-central Italy, one from central–southern Italy, and one corresponding to the sample from southern Italy. The secondary partition, obtained via recursive analysis, largely confirmed the initial grouping, except for the *L. pedemontana* clade from northern-central Italy, which was further subdivided into two distinct clusters. The bPTP analysis recovered thirty-five groups within the genus *Luciola* (see Supplementary Table 3). Focusing on our specimens (Table 3), results were fully congruent with ABGD analysis, identifying seven putative species, coinciding with *L. pedemontana* of Elba clade 1 (support = 0.775), *L. pedemontana* of Elba clade 2 (support = 0.734) and *L. pedemontana* of Elba clade 3 (support = 0.636), *L. pedemontana* of central–southern Italy (support = 0.717), *L. pedemontana* of southern Italy (support = 0.746), and *L. pedemontana* from northern-central Italy split into two distinct clusters, but with low support values (0.587 and 0.493). Finally, the GMYC single-threshold analysis rejected the null hypothesis of a coalescent model (likelihood ratio 16.1042; LR test: < 0.01, significant). It identified seven candidate species within *Luciola* (Supplementary Table 3) but collapsed all our specimens into a single entity comprising *L. pedemontana, L. italica* and *L. lusitanica*. The multiple-threshold analysis also rejected the coalescent model (likelihood ratio 20.24314; LR test: < 0.01, significant), identifying twenty-three species across *Luciola* and five entities in our dataset: two *L. pedemontana* lineages from northern-central Italy, one lineage from central–southern Italy, and one lineage from southern Italy as distinct entities, while the three Elba clades were grouped together as a single entity. In addition, the reference sequences from the Corsica + Sardinia + north Italy lineage, the further Tuscany lineage, and the north-western Italy + France lineage were identified as additional putative species from all the three species delimitation methods (Supplementary Table 3).

**Table 3.** Results from the species delimitation analyses using the ABGD, GMYC and bPTP methods on mtDNA haplotypes (details in the text); different putative species are coded with different letters (A - G).

|  | ABGD |  | GMYC |  | bPTP |
| --- | --- | --- | --- | --- | --- |
|  | Primary | Secondary | Single | Multiple |  |
| *Elba lineage 1* | A | A | A | A1 | A |
| *Elba lineage 2* | B | B | A | A1 | B |
| *Elba lineage 3* | C | C | A | A1 | C |
| *northern-central Italy (Casentino)* | D | D1 | A | A2 | D |
| *northern-central Italy (Tarquinia)* | D | D2 | A | A2 | E |
| *southern Italy* | E | E | A | A3 | F |
| *central–southern Italy* | F | F | A | A4 | G |

## Discussion

Our results confirm the need for more thorough investigation of island biodiversity, uncovering an unexpectedly high level of insect cryptic diversity in an apparently well-studied insular system, the Tuscan Archipelago in the Mediterranean region. Within what was considered a single firefly species, *L. pedemontana*, we found multiple, deeply divergent evolutionary lineages, that exhibit levels of differentiation comparable to, or exceeding, those typically observed among distinct species. Although these findings should be interpreted cautiously due to reliance on a single mitochondrial marker (COI), they provide clear evidence of ongoing evolutionary processes driving lineage diversification. These processes require further investigation and highlight the urgency of their implementation into conservation strategies.

The COI sequence obtained from the single specimen collected on Giglio Island matches *L. italica* with 99.8% identity and represents the first record of this species south of the Po Plain (Fanti, 2022). As this species has a predominantly northern Italian distribution (Fanti, 2022), further investigations are necessary to confirm whether its presence on the island reflects a historically stable population or results from a recent human-mediated colonisation.

On the other hand, the COI sequences from all the other specimens matched with *L. pedemontana*. However, while apparently similar morphologically, phylogenetic analyses recovered 6 strongly differentiated and geographically structured genetic lineages: three from Elba Island and three from the Italian peninsula (Figure 1). The genetic divergence among the Elban clades ranged from 3% to 4.1%, while the divergence between Elban and mainland clades ranged from 3.4% to 10%, values that are above the divergence found between the closely related species *L. italica* and *L. lusitanica* (2.2%), and that exceed the 2–3% threshold commonly applied for delimiting species using the COI barcode region (Hebert et al., 2003; Barrett & Hebert, 2005; Lefébure et al., 2006; Huang et al., 2008; Lukhtanov et al., 2009; Hsu et al., 2013). Species delimitation analyses fully supported the hypothesis of multiple *Luciola* species within our dataset. Both ABGD and bPTP supported the subdivision of *L. pedemontana* into six multiple, distinct species (Table 3). While the results from the two methods were mostly congruent, supporting three species within Elba Island and at least three species across the Italian peninsula, bPTP proposed a further split within the northern peninsular clade, which was also recognised in the ABGD secondary partition. By contrast, GMYC analysis did not return enough resolution, failing to recognise distinct entities in our dataset, even among *L. pedemontana* and the outgroup sequences from *L. italica* and *L. lusitanica*. This discrepancy aligns with other studies reporting GMYC low sensitivity in the presence of uneven sampling and complex demographic histories (Phillips et al., 2019; Magoga et al., 2021), as in this case. When interpreted together, the congruence between distance-based and phylogenetic-based approaches strongly supports the existence of several independent evolutionary lineages within *L. pedemontana* in the study area, a pattern consistent with previous studies documenting marked genetic differentiation within this species across the Italian range (Mori et al., 2025; Nunes et al., 2026).

The spatial pattern of mitochondrial divergence among the *L. pedemontana* lineages observed across the Italian Peninsula reflected the peninsula geographic complexity. Indeed, the main peninsular lineages are differentiated across a north–south axis, with distributions mirroring well-established biogeographic discontinuities, such as those of the northern, central, and southern Apennines. Comparable spatial patterns of genetic divergence have been reported in several other taxa, including insects (Allegrucci et al., 2005; Garzia et al., 2025; Masoni et al., 2025) and vertebrates (Canestrelli et al., 2010, 2013; Senczuk et al., 2017; Chiocchio et al., 2019, 2022). Such geographic patterns have been explained by the effects of Pleistocene climatic oscillations and long-term isolation within distinct refugia on lineage diversification. In this context, the deeper divergence between the northern and the south-central lineages of *L. pedemontana* across the Italian peninsula is consistent with a scenario of prolonged isolation in distinct refugial areas, whereas the further splits within these lineages may indicate more recent divergence within sub-refugia. On the contrary, the lack of substantial differentiation between the firefly populations of central Italy and Ponza supports a scenario of a recent colonisation of this island from the mainland, a pattern also observed in other taxa (Senczuk et al., 2018). However, more samples from other locations are required to clarify the genetic structure and evolutionary history of *Luciola* fireflies across the Italian peninsula.

In contrast, the marked differentiation between the Elban fireflies and both the mainland and Sardinia-Corsica populations, together with the presence of three deeply divergent lineages within Elba Island (*i*.*e*., in an area of no more than 224 km^2^), suggests an ancient colonisation of the island in a more complex biogeographic scenario. Given the limited sampling coverage of the Italian peninsula in our dataset, even in light of the high level of diversification found, we cannot exclude that one or more of these lineages derived from the Italian peninsula. Also, a contribution by source populations from Corsica cannot be excluded, given documented Corsica–Elba affinities in many taxa (see *e*.*g*. Cianchi et al., 2003). However, the lower genetic divergence observed between the Elban and peninsular lineages, compared with that between the Elban and Sardinia–Corsica lineages, points to a closer biogeographic and evolutionary affinity of Elban lineages with the Italian Peninsula than with the Sardinia–Corsica system. In the current state of knowledge, and pending broader regional investigation on firefly distribution and genetic structure, the most parsimonious interpretation is that the three Elban lineages represent insular endemic evolutionary units.

Notably, the spatial distribution of the Elban lineages indicates a weak, but consistent east–west structure, with partial sympatry in the central area and predominance of distinct clades in the northeastern and southeastern sectors. Although these boundaries are diffuse, this spatial pattern is consistent with the island’s heterogeneous topography and fine-scale environmental gradients in maintaining partial lineage isolation. Because of the acknowledged limitations of mtDNA markers in describing fine-scale patterns of population structure, these results should be interpreted with caution and recommend further investigation using multilocus genomic datasets (Catalán et al., 2024). Finally, cytoplasmic endosymbionts such as *Wolbachia* may bias mitochondrial variation by generating rapid cytoplasmic sweeps leading to instances of cytonuclear discordance (Hurst & Jiggins, 2005; Smith et al., 2012). Because *Wolbachia* infection status was not assessed in our dataset, we cannot rule out the possibility that these processes contributed to the observed geographic patterns.

## Conclusions

This study provides the first molecular perspective on *Luciola* diversity within the Mediterranean region. In particular, our results support the that Mediterranean islands harbour ecological processes and historical biogeographic dynamics that promote evolutionary diversification from nearby islands and mainland regions. From a conservation perspective, the presence of deeply divergent lineages confined to a small and isolated area and the awareness of the many factors that are causing widespread declines of firefly populations, such as habitat loss, artificial light at night, and pesticide use (Lewis et al., 2020, 2024), identify the Tuscan Archipelago as a priority area for firefly conservation. Moreover, the high level of diversification found outlines the Tuscan archipelago as a local biodiversity hotspot for fireflies, a role already documented for several other taxa with island-restricted lineages, including plants (Coppi et al., 2014; Foggi et al., 2015; Carta et al., 2018), invertebrates (Fattorini, 2009; Barbato et al., 2018; Forbicioni et al., 2019) and vertebrates (Ancillotto et al., 2026). The loss of such biological diversity would entail the erosion of unique evolutionary history within an already vulnerable Mediterranean island system (Marescalchi et al. 2025).

## Acknowledgment

We would like to thank Armando Macali, Marco Rinaldi, Francesco Cappuccio, Andrea Caliendo, Edoardo Tatti, Chiara Tatti, Dario Boldrini, Giandomenico Catanese, Michael Piras, Davide Mancani, and Federico Matafirri for their help on fieldwork, and Michela Paoletti for her valid support during laboratory procedures.

## Data availability

The dataset generated during the current study is available in the GenBank repository (accession numbers listed in Supplementary materials).

## Author contributions

**E. Serafini**: data curation (equal); investigation (lead); writing – original draft (lead); formal analysis (lead); writing – review and editing (equal); visualization (lead). **A. Chiocchio**: data curation (equal); investigation (equal); writing – original draft (lead); formal analysis (lead); writing – review and editing (equal); visualization (equal); supervision (lead). **L. Forbicioni**: data curation (equal); investigation (equal); writing – review and editing (equal). **A. Lagrotteria:** data curation (equal); writing – review and editing (equal). **A. Viviano:** data curation (equal); writing – review and editing (equal). **E. Mori:** data curation (lead); writing – original draft (equal); review and editing (equal). **L.Ancillotto:** data curation (equal); writing – review and editing (equal). **R. Bisconti:** funding acquisition (equal); review and editing (equal). **D. Canestrelli:** conceptualization (lead); funding acquisition (lead); supervision (equal); review and editing (equal).

